# Transmission of vibrations in buzz-pollinated plant species with disparate floral morphologies

**DOI:** 10.1101/2021.04.16.440147

**Authors:** Lucy Nevard, Avery L. Russell, Karl Foord, Mario Vallejo-Marin

## Abstract

In buzz-pollinated plants, bees apply vibrations produced by their thoracic muscles to the flower, causing pollen release from anthers, often through small apical pores. During floral buzzing, bees grasp one or more anthers with their mandibles, and vibrations are transmitted to the focal anther(s), adjacent anthers, and the whole flower. Because pollen release depends on the vibrations experienced by the anther, the transmission of vibrations through flowers with different morphologies may determine patterns of release, affecting both bee foraging and plant fitness. Anther morphology and intra-floral arrangement varies widely among buzz-pollinated plants. Here, we compare the transmission of vibrations among focal and non-focal anthers in four species with contrasting anther morphologies: *Cyclamen persicum* (Primulaceae), *Exacum affine* (Gentianaceae), *Solanum dulcamara* and *S. houstonii* (Solanaceae). We used a mechanical transducer to apply bee-like artificial vibrations to focal anthers, and simultaneously measured the vibration frequency and displacement amplitude at the tips of focal and non-focal anthers using high-speed video analysis (6,000 frames per second). In flowers in which anthers are tightly held together (*C. persicum* and *S. dulcamara*), vibrations in focal and non-focal anthers are indistinguishable in both frequency and displacement amplitude. In contrast, flowers with loosely arranged anthers (*E. affine*) including those in which stamens are morphologically differentiated within the same flower (heterantherous *S. houstonii*), show the same frequency but higher displacement amplitude in non-focal anthers compared to focal anthers. Our results suggest that stamen arrangement affects vibration transmission with potential consequences for pollen release and bee behaviour.

## Introduction

Insects use substrate-borne vibrations in a range of ecological contexts, including conspecific communication and the detection of prey and predators (Cocroft and Rodríguez, 2005; Mortimer, 2017). These vibrations are often produced and detected on plant material, and the physical properties of the plant substrate, such as stem stiffness or leaf thickness, often affect vibration propagation (Cocroft et al., 2006; Kollasch et al., 2020; Velilla et al., 2020). Approximately 6-8% of angiosperms are buzz-pollinated, relying on substrate-borne vibrations (floral buzzing), typically produced by bees, to release pollen from flowers with specialised morphologies (Buchmann, 1983; Vallejo-Marín, 2019). While buzz pollination is a widespread plant-insect interaction common in agricultural and natural ecosystems, its biomechanical aspects remain understudied compared to other insect vibrations.

More than half of all bee species can buzz to collect pollen, and the behaviour is thought to have evolved approximately 45 times within bees (Anthophila)(Cardinal et al., 2018). During buzz pollination, the bee typically clutches the anthers with its mandibles and produces thoracic vibrations using the indirect flight muscles (Buchmann, 1983; King and Buchmann, 2003). These vibrations are transmitted to the flower, triggering pollen release. Most buzz-pollinated flowers have tubular anthers that dehisce only via small apical pores or slits, i.e., poricidal anthers, through which small, dry pollen grains are released during floral vibrations (Harder and Barclay, 1994). Moreover, some species with longitudinally dehiscent anthers have evolved floral morphologies which also rely on floral buzzing for pollen release. For example, the modified corolla of some *Pedicularis* species encloses the anthers in a tube, which thus functions analogously to an individual poricidal anther (Corbet and Huang, 2014). Furthermore, many species with non-poricidal anthers and apparently accessible pollen, e.g., *Rosa* or *Begonia* species, are often buzzed by bees, presumably maximizing pollen collection (Buchmann, 1985; Russell et al., 2020). The interaction between flower and vibrating bee is thus very widespread, emphasising the importance of studying floral vibrations in detail across plant lineages.

Similar to the study of vibrations used for insect communication, the functional study of floral vibrations can be divided into three major stages: (1) the production of vibrations by the bee, (2) the propagation of these vibrations through the bee-flower coupled system, and (3) the effect of vibrations on pollen release (Oberst et al., 2019; Vallejo-Marin 2019). Most work to date has focused on (1) bee buzzing behaviour and/or (3) pollen release. Bees produce floral vibrations which vary in duration, frequency and amplitude, the primary components with which vibrations can be described. Vibration amplitude, whether measured as displacement, velocity or acceleration, has a significant and positive effect on pollen release: higher amplitude vibrations release more pollen (De Luca et al. 2013; Kemp and Vallejo-Marin, 2020; Rosi-Denadai et al., 2020). In contrast, the effect of vibration frequency on pollen release appears to be weaker within the natural range of bee buzzes (~100 - 400 Hz; De Luca et al., 2019; De Luca and Vallejo-Marín, 2013; Rosi-Denadai et al., 2020), although vibrations at much higher frequencies than those produced by bees do result in the release of more pollen (Arceo-Gómez et al., 2011; Harder and Barclay, 1994).

Buzz-pollinated flowers are morphologically diverse, yet the intra-floral transmission of vibrations across a range of species has been rarely investigated. The structure of the androecium, e.g., the spatial arrangement of the anthers, is likely to affect transmission of vibrations. Many taxa with poricidal anthers have converged on a floral morphology in which equally sized anthers are held tightly together forming an anther cone as in *Solanum dulcamara* L. and *S. lycopersicum* L. (Glover et al., 2004). Interestingly, this anther cone morphology has evolved independently in many other groups of flowering plants (Faegri, 1986; Glover et al., 2004; Vallejo-Marín et al., 2010). During floral buzzing, bees often use their mandibles to hold only a subset of the anthers in the flower (Papaj et al., 2017). In species with tightly arranged anthers, bee vibrations applied to one or a few anthers are likely to be effectively transmitted to the rest of the anther cone. In contrast, in buzz-pollinated species with anthers presented more loosely (e.g., most *Melastomataceae*, *Solanum elaeagnifolium* Cav., *S. sisymbriifolium* Lam.), applying vibrations to a subset of focal anthers might limit transmission to non-focal anthers in the same flower.

This potential difference in vibration transmission between focal and non-focal anthers is perhaps best exemplified in heterantherous species, in which two or more morphologically distinct sets of anthers occur in the same flower (Vallejo-Marín et al., 2010). In some heterantherous species the two anther sets perform different functions, with long anthers contributing disproportionately to fertilisation (pollinating anthers) and short anthers (feeding anthers) being the focus of attention of buzz-pollinating bees (Luo et al., 2008; Müller, 1881; Vallejo-Marin et al., 2009). A recent study has shown that pollinating and feeding anthers of heterantherous *Solanum* have different natural frequencies, likely as a result of differences in biomechanical properties including size and shape (Nunes et al., 2020). Despite the potential for differences in anther and floral characteristics, such as those described above, to affect the transmission of vibrations in buzz-pollinated flowers, few studies have explicitly compared floral vibrations across disparate floral morphologies.

Here, we used a mechanical transducer to apply bee-like artificial vibrations to focal anthers, simultaneously measuring the vibration frequency and displacement amplitude at the tips of focal and non-focal anthers of the same flower in two axes using high-speed video analysis (6,000 frames per second). We used four buzz-pollinated species with contrasting floral and poricidal anther morphologies: *Cyclamen persicum* Mill. (Primulaceae), *Exacum affine* Balf. ex Regel (Gentianaceae), *Solanum dulcamara* and *S. houstonii* Dunal (Solanaceae). The arrangement of anthers within these flowers varies from a tight cone (*S. dulcamara*) to a loose, heterantherous assemblage (*S. houstonii*). We ask the following questions: i) Does the fundamental frequency of vibrations change between focal and non-focal anther in these flowers? ii) Does vibration amplitude (measured as displacement amplitude) change between focal and non-focal anthers? iii) Do vibration characteristics depend on plant species and/or the characteristics of the applied vibration? Based on previous work suggesting conservation of frequency properties during buzz pollination (De Luca et al., 2020; Pritchard and Vallejo-Marín, 2020), but changes in amplitude as vibrations travel through the flower (Arroyo-Correa et al., 2019; King and Buchmann, 1996), we predict that vibration amplitude, but not frequency, will be transmitted from focal to non-focal anthers less faithfully in flowers with looser anther arrangements. High-speed video requires no physical interference with the system and is an alternative to other non-contact methods to study vibrations across complex structures such as laser scanners. Our study allows us to quantify and compare the transmission of floral vibrations in flowers with disparate morphologies and may be useful for future work on the function and evolution of different floral morphologies among buzz-pollinated plants.

## Materials and Methods

### Study system and plant material

We studied flowers of four species from three families: *Cyclamen persicum* (Primulaceae), *Exacum affine* (Gentianaceae), *Solanum dulcamara* and *S. houstonii* (Solanaceae). These species have contrasting stamen morphologies and are nectarless, offering only pollen as a reward to floral visitors (Fig. 1B-F). *Cyclamen persicum* flowers have poricidal anthers fused by the filaments into a symmetrical conical shape (Schwartz-Tzachor et al., 2006). Together with other *Cyclamen* species, *C. persicum* was historically presumed to be buzz-pollinated, based on the presence of poricidal anthers. However, buzz-pollinating visitors are rarely observed in wild populations and its main pollinators are often moths, hoverflies and small bees (Schwartz-Tzachor et al., 2006). *Exacum affine* has freely moving, slightly curved, poricidal anthers and is primarily buzz-pollinated (Endress, 2012; Russell et al., 2015). *Solanum dulcamara* has poricidal anthers which are fused along their length into a tight cone (Glover et al., 2004). *Solanum houstonii* has freely moving, curved, poricidal anthers and is heterantherous: it has two short anthers (feeding anthers) presumed to be mainly involved in attracting and rewarding pollinators, and three longer, S-shaped anthers (pollinating anthers) presumed to contribute disproportionally to pollination (Papaj et al., 2017). Both *Solanum* species are buzz-pollinated by bees of diverse sizes and morphologies (Carbonell, 2019; Free, 1970; Glover et al., 2004; Macior, 1964; Papaj et al., 2017; L.N., personal observation) (Fig. 1).

**Figure 1.**
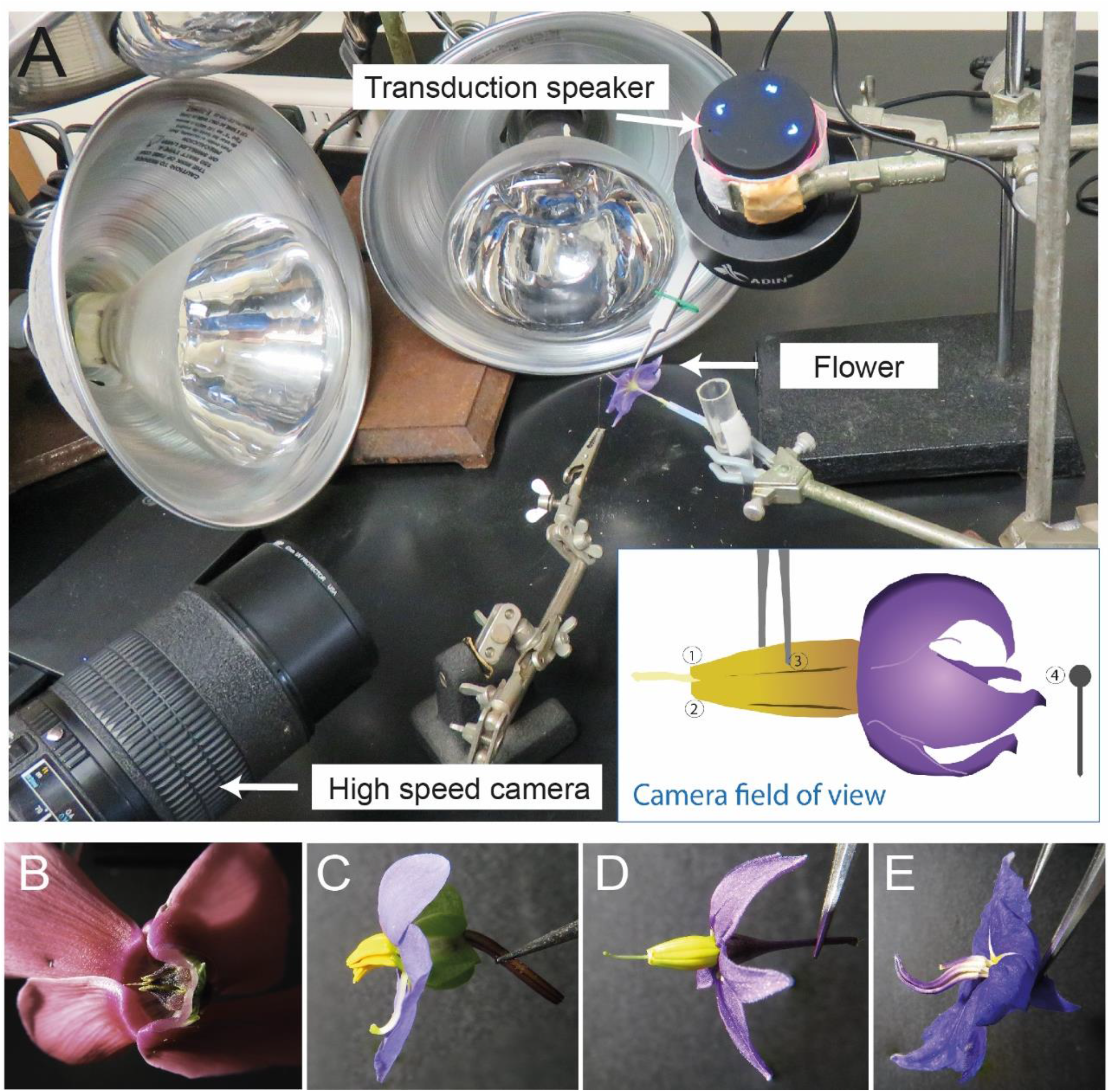
A: Photo of artificial vibration playback with inset diagram of camera field of view. 1: Focal anther tip; 2: non-focal anther tip; 3: forceps tip; 4: insect pin for calibration. Example flower is *S. dulcamara*. B: *Cyclamen persicum*; C: *Exacum affine*; D: *Solanum dulcamara*; E: *Solanum houstonii*.

Plants were purchased as full-grown plants or grown from seeds or cuttings in university greenhouses in Tucson, AZ and Pittsburgh, PA. *Cyclamen persicum* plants were sourced from Lowe’s Home Improvement. *Exacum affine* plants of three varieties (Champion Blue, Royal Blue, Little Champ Blue) were sourced from the wholesaler Fred C. Gloeckner & Co. *Solanum dulcamara* cuttings were collected from wild populations in Pittsburgh, PA. *Solanum houstonii* seeds were sourced from the Sonoran Desert Museum, Tucson, AZ; originally collected from wild populations in Mexico.

### Synthesis and playback of vibration signals

The experiment consisted of generating synthetic vibrations and applying them to individual flowers using a vibration transducer mechanical system, to analyse the vibrational properties of different parts of the flower. Flowers used in experiments were as fresh as possible, usually newly opened on the plant each morning of the experiment. Synthetic vibrations consisted of 1 s pure tone signals of fixed amplitude with a 10 ms fade in and 10ms fade out. We conducted two sets of experiments. In the first set, we varied relative amplitude of the signal, while keeping frequency constant. Signals were generated by creating a sine wave with frequency of 350 Hz, using the Tone function in Audacity ver. 2.1.0 (http://audacityteam.org/) and saved as a single-channel audio file (WAV) at 44.1 kHz sampling rate. We used four relative amplitude levels: (in dB): −15, −10, −5 and 0. The absolute displacement amplitude of the vibrations applied to the flower in each of these treatments was calculated using the observed displacement of the forceps tips (see *Digitising Video Files and Time Series Analysis* section). For each amplitude, we conducted 2-3 playback replicates per species in each of two species (*Exacum affine* and *Solanum houstonii*). In the second experiment we generated signals as above with constant amplitude (0dB) but with different individual frequencies (150, 200, 250, 300, 350, 400, 450, and 500 Hz). The frequency values we used reflect the range of frequencies recorded from bees vibrating on buzz-pollinated flowers (De Luca et al., 2019; De Luca and Vallejo-Marín, 2013; Vallejo-Marín and Vallejo, 2021). For each frequency, we conducted 3-4 playback replicates for each of the four species studied, depending on flower availability.

We played back each vibration signal using a Zoom H2 audiorecorder (Zoom Corporation; Tokyo, Japan) connected to a vibration speaker (Adin S8BT 26W). The output volume of the Zoom H2 and vibration speaker was kept constant, except as noted in the Results section. The vibration speaker was modified as described in Brito et al. (Brito et al., 2020) to transduce the vibrations to the flower by fixing a metal rod to the vibrating plate of the speaker and attaching a pair of very fine tipped forceps (Fine Science Tools, Dumont #5 Biology Tip Inox Forceps) to the end of the rod. The forceps were used to hold 1-2 anthers (the short anthers in the case of *Solanum houstonii*) (see Fig. 1A for setup). Individual flowers were placed in floral water tubes, with the stamen’s long axis parallel to the ground, i.e., flowers were kept horizontal to the ground as they would be perceived by a pollinator directly approaching the centre of the flower. The forceps were clamped at approximately the same position on the anthers for each trial. A fresh flower was used for each replicate such that each flower was vibrated only once and we collected data sequentially for each amplitude level or frequency before moving to the next set of replicates to control for effects of time of day on vibration characteristics.

### High speed digital imaging

To analyse the vibration of different parts of the flower simultaneously, we used high-speed digital imaging, which allowed us to simultaneously track the movement of captured objects along two dimensions at different locations of the image frame. We recorded the vibrating flowers at 6,000 frames per second (fps; 1280 x 512 pixels) against a black background using a FASTCAM SA-8 camera (Photron, San Diego, California USA) and halogen bulbs for illumination. Recording started before the vibration playback began and captured the whole 1 second vibration. An entomological pin (size 1) was kept in shot for videos, to enable size calibration and consistency in video tracking output across different videos (Fig 1).

### Digitising video files and time-series analysis

All video footage was analysed in two dimensions using the *DLTdv7* digitising tool (Hedrick, 2008) in MATLAB 9.6 (R2019a; MathWorks Inc). Recordings were 730 ms long on average. This digitising tool allows point tracking in high-speed video footage (Varennes et al., 2019), and we used it to generate a time series of *x-y* coordinates for each tracked point. For each video, we simultaneously tracked three points through time to extract vibrational information measured as displacement: (1) The tip of the forceps, hereafter *control*. This allowed us to empirically obtain frequency and displacement amplitude of the input vibrations transduced to the flowers, and to account for variation in volume playback introduced during the experiment (Fig. S1). (2) The tip of the anther held by the forceps, hereafter the *focal anther*. (3) The tip of the anther furthest away from the focal anther, hereafter the *non-focal anther*. In a few cases, it was not possible to track all three points for each sample due to obstruction of the control point by other parts of the flower or due to low light. When the control point could not be tracked, the recording was not used in downstream analyses.

The x-y time series data was analysed using the *seewave* package (Sueur et al., 2008) in R ver. 4.0.2 (R Core Development Team, 2020). Displacement values (calibrated to mm, using the insect pin described previously as a reference for size) were calculated for x- and y-axes, by zero-centring the data. These x-y displacements were used to obtain an overall measure of displacement magnitude defined as (x-displacement^2^ + y-displacement^2^)^1/2^. We used a high pass filter of 80Hz using the *fir* function (Hanning window, window length = 512 samples). For each digitised recording, a section of 100 ms in the middle of each time series was selected, where the vibration was more stable (approximately from 0.3 seconds to 0.4 seconds for every sample). Digitised samples which were too short were removed from the dataset. From these 100 ms sections, we computed the frequency spectra using the function *spec* (using power spectral density) and calculated the dominant or peak frequency using the function *fpeaks* (nmax =1). We also estimated peak displacement amplitude (D_P_), peak-to-peak displacement (D_P-P_), and Root Mean Squared (D_RMS_) displacement using the functions *max* (on absolute values), *max – min*, and *rms*, respectively.

### Statistical analysis

We evaluated the correlation between the different measurements of displacement amplitude (D_P_, D_P-P_, D_RMS_), and between displacement in the x-, y-axis and v-vector using Pearson moment correlations. We assessed the association between the characteristics of the input vibrations applied by the forceps (peak frequency and D_RMS_) and those measured at the anther tips using linear models fitted with the function *lm*. In these models, vibration peak frequency or D_RMS_ were used as the response variable, and input vibration (at the forceps tip), anther type (non-focal and focal anthers) and species as the explanatory variables. For each model, diagnostics were produced using the package DHARMa (Hartig, 2019). For those which showed significant outliers, models were re-created without these data points. The statistical significance of effects remained similar and therefore we kept the full data set for the final analysis. Statistical significance of the main effects and their interactions were assessed using Type III sums of squares using the package *car* (Weisberg and Fox, 2011). Model predictions were plotted using *plot_model* (type=pred) in the package *sjPlot* (Ludecke, 2021). All statistical analysis was performed in R 4.0.2 (R Core Development Team, 2020).

## Results

### Frequency of anther vibrations

The peak frequencies measured in the x- and y-axis were highly correlated across all samples. (Pearson’s correlation *r*: 0.98, df: 266, p < .001) (Fig. 2A). Peak frequency across anthers and plant species ranged from 150 to 529Hz (Fig. S3). Forceps peak frequency was the only significant predictor of anther peak frequency in our linear model (p<0.001, Table 1A, Fig. S2) and we found no effect of either anther type or plant species (i.e., anther arrangement) on the peak frequency of vibrations measured at the tips of anthers (p > 0.05, Table 1A, Fig. S2). In other words, the peak frequency did not change as vibrations were transmitted through the flowers from the forceps. The overall frequency spectra were also similar between species and anther type, with very few harmonics in any of the vibrations (Fig. 3).

**Figure 2.**
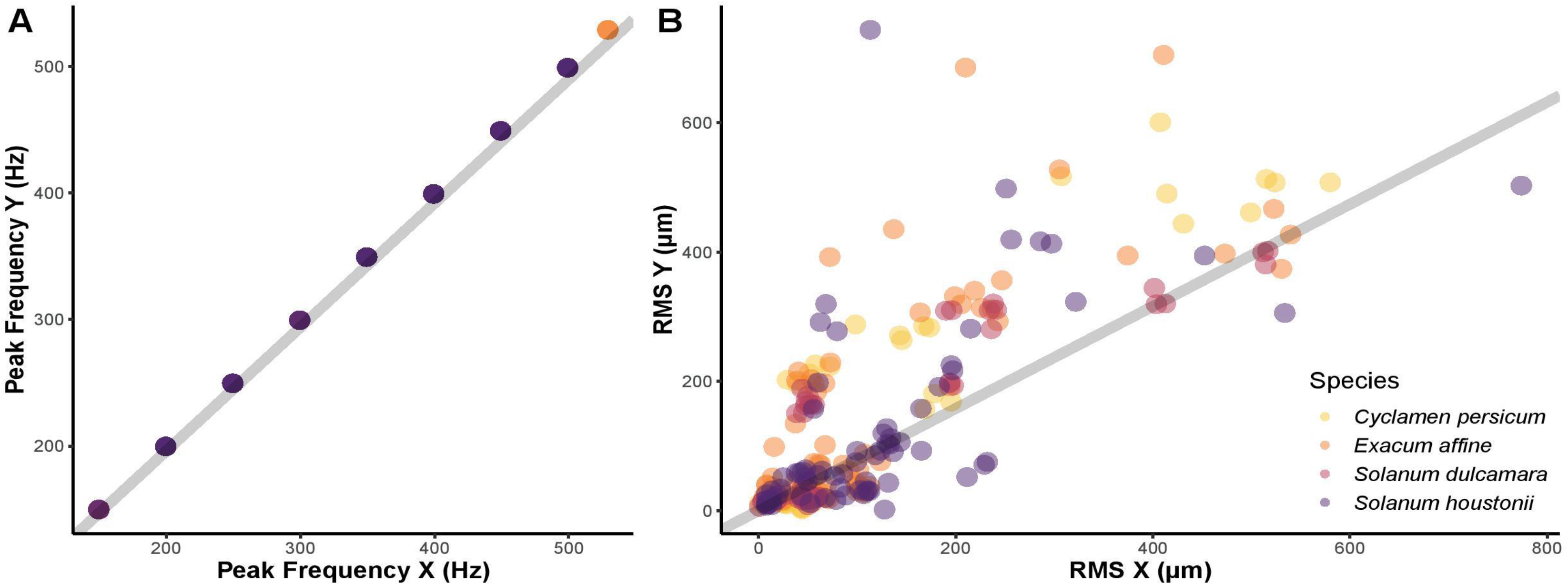
Correlation of frequency and amplitude (D_RMS_) variables in x and y axes across all samples (Pearson’s correlation coefficient). Grey lines indicate correlation. A: peak frequency (x) v peak frequency (y) (r = 0.98). B: D_RMS_ (x) v D_RMS_ (y) (r = 0.79).

**Table 1.** Linear models fitted with peak frequency (Hz) or D_RMS_ respectively as response, and forceps peak frequency or D_RMS_, anther type and species as fixed effects. \**P*-value of fixed effect in linear model. \*\**P*-value calculated using Type III sums of squares. Sample size is 150 for both models.

|  | Estimate | Std. error | *P*-value* | *P*-value** |
| --- | --- | --- | --- | --- |
| **A. Peak frequency (Hz)** |  |  |  |  |
| Forceps peak frequency | 1.000e+00 | 1.552e-16 | **<0.001** |  |
| Anther (non-focal) | 3.459e-14 | 3.077e-14 | 0.263 |  |
| Species |  |  |  |  |
| (*Exacum affine*) | 6.634e-14 | 4.279e-14 | 0.123 |  |
| (*Solanum dulcamara*) | 7.741e-14 | 4.711e-14 | 0.103 |  |
| (*Solanum houstonii*) | 6.576e-14 | 4.446e-14 | 0.141 |  |
| **B. Displacement amplitude D<sub>RMS</sub> (μm)** |  |  |  |  |
| Forceps D<sub>RMS</sub> | 1.012 | 0.045 | **<0.001** | **<0.001** |
| Anther (Non-focal) | -16.825 | 9.464 | 0.078 | 0.078 |
| Species |  |  |  | 0.13 |
| (*Exacum affine*) | -2.68 | 7.917 | 0.735 |  |
| (*Solanum dulcamara*) | -2.969 | 8.775 | 0.736 |  |
| (*Solanum houstonii*) | 12.441 | 8.05 | 0.124 |  |
| Forceps D<sub>RMS</sub>: Anther (non-focal) | 0.202 | 0.064 | **0.002** | **0.002** |
| Anther (non-focal): Species |  |  |  | **<0.001** |
| Anther: (*Exacum affine*) | 44.754 | 11.306 | **<0.001** |  |
| Anther: (*Solanum dulcamara*) | 1.31 | 12.303 | 0.915 |  |
| Anther: (*Solanum houstonii*) | 40.533 | 11.792 | **<0.001** |  |

**Figure 3.**
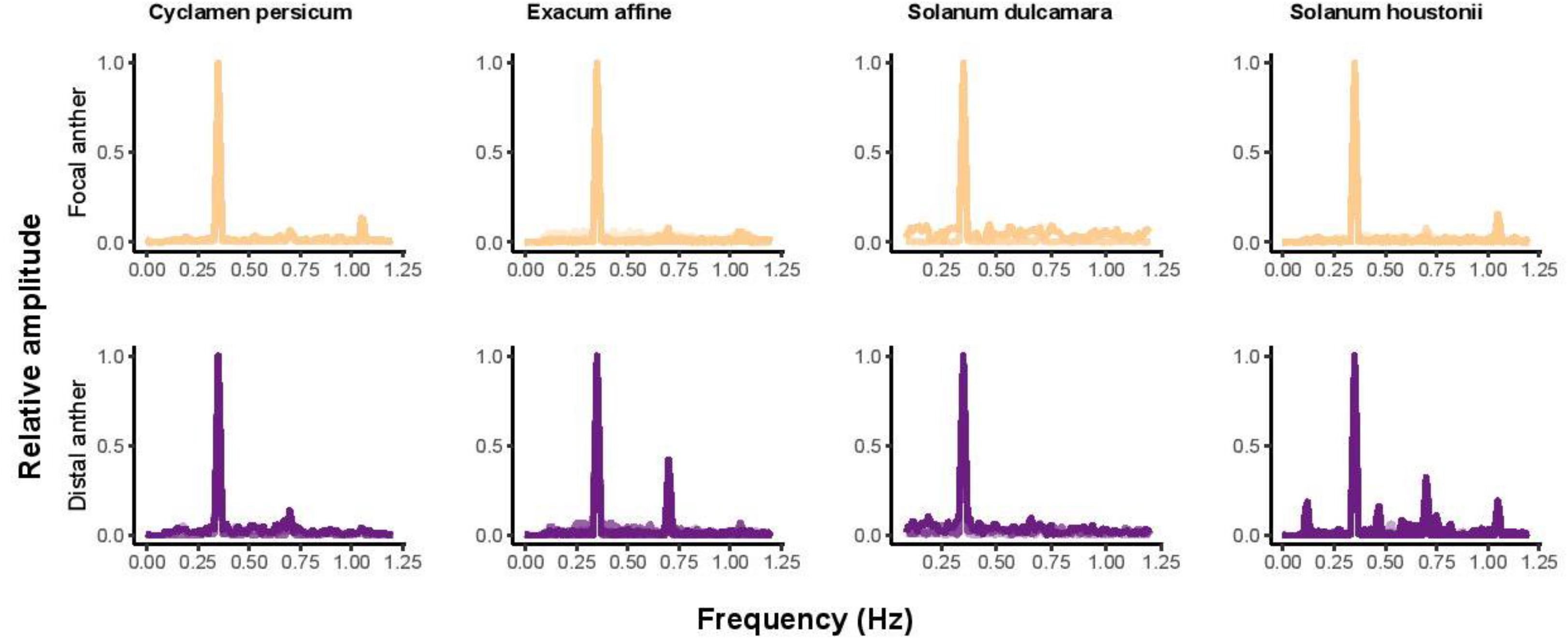
Number of peaks (above 0.4 of total amplitude) in frequency spectra against input frequency (Hz) for focal and distal anther of four plant species, at an input frequency of 350Hz.

### Amplitude of anther vibrations

All three measures of displacement amplitude differed slightly between the x- and y-axis across all samples, including in the forceps (Table 2). The average amplitude was higher in the y-axis, particularly in D_P_ and D_P-P_ (Table 2). Axes were nonetheless strongly correlated for all measures of amplitude: D_P_ (r: 0.81, df: 266, p < 0.001); D_P-P_ (r:0.82, df: 266, p <0.001); D_RMS_ (r: 0.79, df:266, p < 0.001) (Fig. 2B for D_RMS_ correlations). Therefore, we used the vector magnitude (see Methods for details) for downstream analysis on amplitude, to capture variation in displacement in both x and y axes.

**Table 2.**
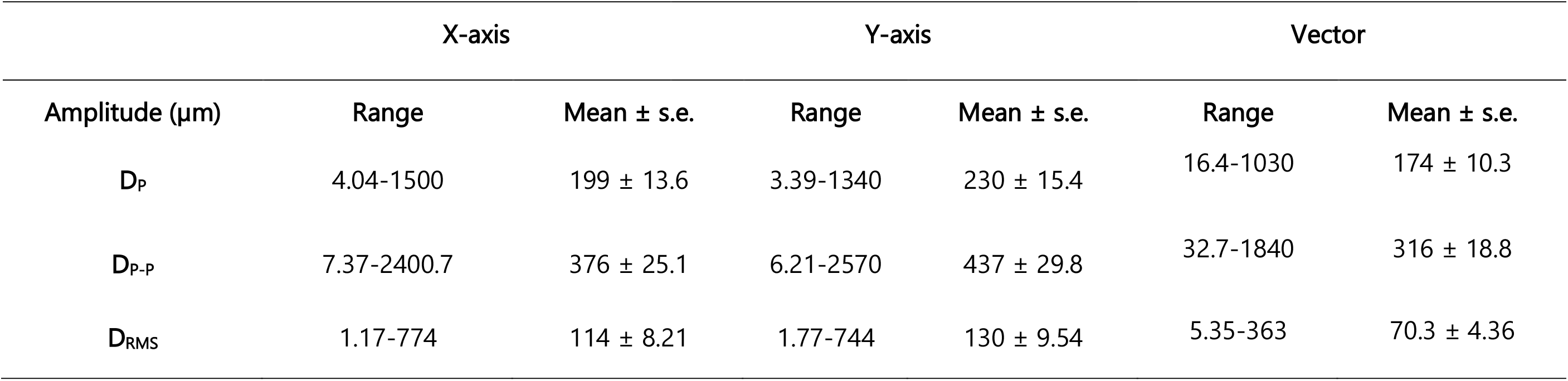
Ranges and means, across all samples, of three measures of displacement amplitude (μm), D_P_, D_P-P_ and D_RMS_, in the x and y axes, and the vector taken from these axes (see Methods).

We extracted three measures of displacement amplitude: D_P_, D_P-P_, and D_RMS_. D_P_ across anther types and species ranged from 16.4μm to 1030μm (mean 195), D_P-P_ ranges from 39.3 to 1840 μm (mean 353), D_RMS_ ranged from 6.94 to 363μm (mean 77.8). The highest displacements for all measures were from vibrations in the non-focal anther of *S. houstonii*, and the lowest were from the non-focal anther of *C. persicum*. All three measures of displacements were strongly correlated across all trials: D_P_ and D_P-P_ (r: 1, df: 179, p < 0.001); D_P_ and D_RMS_ (r: 0.98, df: 179, p < 0.001); D_P-P_ and D_RMS_ (r: 0.99, df:179, p < 0.001). D_RMS_ was used for all further amplitude analysis.

We found a significant interaction between anther type and input D_RMS_ (measured at the forceps) on anther displacement (vector D_RMS_), with displacement in non-focal anthers generally increasing more rapidly with input amplitude than in focal anthers (Table 1B, Fig. 4). We also found a significant interaction effect between anther type and plant species on the displacement amplitude (vector D_RMS_) of vibrations (p < 0.001, Table 1B, Fig. 4), with higher displacements in the non-focal anthers of *E. affine* (coefficient = 42.87) and *S. houstonii* (coefficient = 46.11) compared to focal anthers of *C. persicum* (Table 1b, Fig 5). Separate analyses of the x and y axes both showed significant interactions between anther type and plant species (p<0.005) (Figs S4 and S5; Tables S1 and S2). When we calculated the disparity in D_RMS_ (vector) between the forceps and the anther, the mean difference across both anther types in *C. persicum* and *S. dulcamara* was close to zero (Table 3). In contrast, the mean differences for the non-focal anthers of *E. affine* and *S. houstonii* were 36.6μm and 55.1μm respectively, and for the focal anther of *S. houstonii* it was 13.8μm (Table 3).

**Figure 4.**
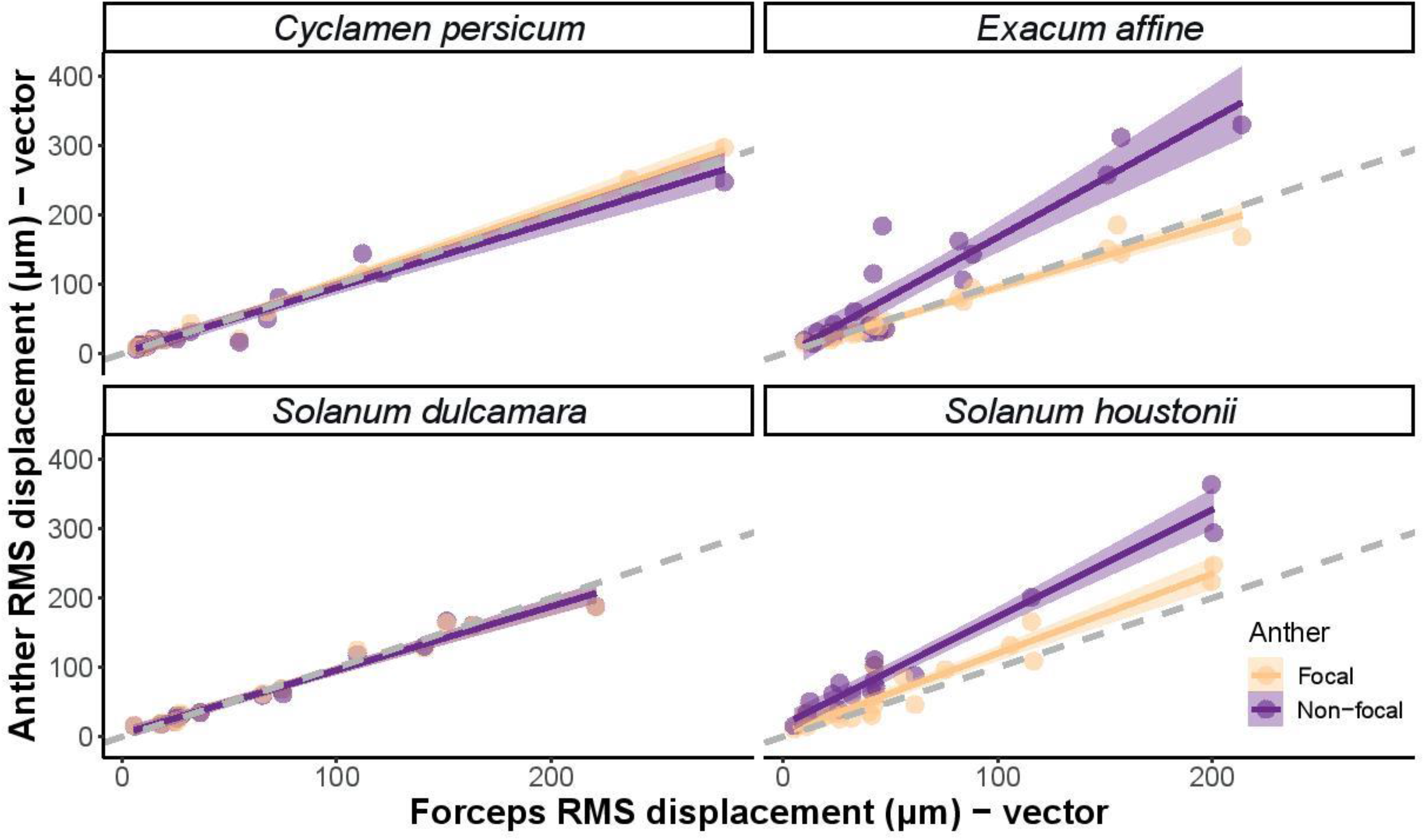
Linear model estimates and data points for measured vector RMS displacement (μm) of focal and distal anther against forceps RMS displacement (μm) in four plant species. Grey dashed line indicates a linear relationship with slope=1.

**Table 3.**
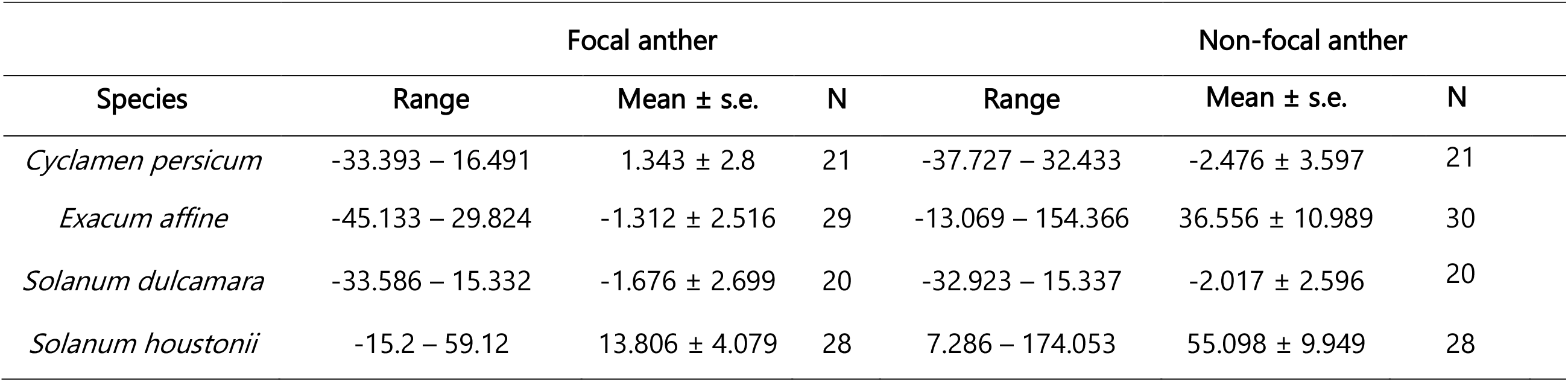
Difference in displacement amplitude in μm (Anther D_RMS_ – Forceps D_RMS_) between forceps and anther for each anther type and species across samples. Values are for the vector calculated from the axes (see Methods for details).

## Discussion

Our study suggests that the arrangement of poricidal anthers affects the transmission of vibrations between anthers. We found that vibrations are transmitted similarly, in both frequency and amplitude, across focal and non-focal anthers in species with stamens partially or totally fused to form a cone (*S. dulcamara* and *C. persicum*). In contrast, species in which individual stamens can move freely (*E. affine* and *S. houstonii*) showed identical frequency but higher vibration amplitudes at the tip of non-focal anthers compared to the focal anthers where vibrations were applied. Overall, the highest displacements occurred in the long anthers of the heterantherous *S. houstonii*. Our work shows that floral morphology, including the functional fusion of stamens into an anther cone, affects the transmission of vibrations applied to a subset of anthers. Because buzz-pollinating bees usually grasp only one or few anthers during buzz pollination and because pollen release is a function of vibration amplitude, our results suggest that floral morphology will be an important determinant of the functional consequences of the applied vibrations, including how bee vibrations might translate into pollen released by flowers with different architectures.

### Transmission of vibrations

The peak frequency of artificial vibrations did not change as they were transmitted through flowers, regardless of flower type or vibration characteristics. This result aligns with Brito et al. (2020) who also found that artificial vibration peak frequency is conserved throughout the heterantherous flowers of *S. rostratum* Dunal, both at anther tips and petals. Although some plant substrates can act as frequency filters (Cocroft et al., 2006), frequency is not altered over the short distances involved in vibration transmission during buzz-pollination interactions (De Luca and Vallejo-Marín, 2013). Although the natural frequency of anthers is affected by their morphology and organisation within the flower (Nunes et al., 2020), the frequency of vibrations has limited effects on pollen release in buzz-pollinated flowers, suggesting that resonance plays a minor role within the range of frequencies produced by most bees (100 to 400Hz) (De Luca and Vallejo-Marín, 2013; Nunes et al., 2020; Rosi-Denadai et al., 2020).

In contrast, we found that the amplitudes of artificial vibrations were differentially altered as they travelled through the two types of buzz-pollinated flowers. In the flowers with more loosely arranged androecia, *E. affine* and *S. houstonii*, vibrations at the tip of the non-focal anther had generally higher displacement amplitude than those observed in the tip of anthers being vibrated. This effect was strongest in the heterantherous *S. houstonii*, where in some cases, displacement was doubled between input and the non-focal anther. In *S. rostratum*, velocity amplitude from the vibration source to the anther tips of both feeding and pollinating anthers increases up to four-fold when vibrations were applied at the base of the flower (Brito et al., 2020). Stamens can be thought of as a complex cantilever beam (a structure with one fixed end and one free end) (King, 1993). Vibration displacement amplitude at the tip of the stamen should be partly a function of stamen’s length, second moment area, Young’s modulus and mass. We thus expect longer stamens to show generally higher displacements at the tip than shorter stamens. Stamen length differences may help explain the difference in vibration amplitude between the short anthers of *E. affine* and the long pollinating anthers of *S. houstonii*. However, stamen material properties and morphology are likely to affect important parameters determining their vibrational properties, including their second moment area and Young’s modulus, and predictions based on length alone might not capture the behaviour of real stamens (Vogel, 2013). Previous empirical work shows that amplitude has a significant positive effect on pollen release (De Luca et al., 2013; Harder and Barclay, 1994; Kemp and Vallejo-Marin, 2020) with increased anther acceleration causing pollen grains to gain in energy and escape through the pores at a higher rate (King and Lengoc, 1993). Clearly more work in this area is needed, including both empirical and modelling studies of the vibrational properties of stamens incorporating the complexity of the forms and material properties of stamens.

Unlike the higher vibration amplitude observed in non-focal anthers of species with loosely held stamens, species in which anthers are tightly held together forming anther cones (*C. persicum* and *S. dulcamara*) showed a negligible change in amplitude between focal and non-focal anthers. The functionally cohesive androecium in these species appears to faithfully transmit vibrations across the anther cone. The uniformity of the amplitude and frequency of vibrations observed in species with anther cones might have implications for patterns of pollen release during buzz pollination. Species with anther cones may show a more uniform rate of pollen release from each anther when vibrated, compared to the more heterogenous range of vibrations experienced by individual anthers of species in which anthers move more freely. Anther cones have evolved in a variety of taxa with buzz-pollinated flowers including species in the families Solanaceae, Primulaceae, Rubiaceae, Gesneriaceae, and Ericaceae (Glover et al., 2004; Harder and Barclay, 1994; Puff et al., 1995; Schwartz-Tzachor et al., 2006) providing excellent opportunities to compare the functional significance of convergent floral morphologies. The same putative uniform pollen release may also occur when non-poricidal anthers are enclosed in a corolla and flowers are buzz-pollinated, as seen in some *Pedicularis* species (Corbet and Huang, 2014; Tong et al., 2019). Our study did not investigate pollen release patterns in different types of flowers and further work quantifying vibratory pollen release in flowers with disparate morphologies across taxonomic groups could help establish the functional consequences, if any, of different androecium architectures.

We suggest that the differences in vibration transmission we see in this study are largely due to differences in anther arrangement in our chosen flower types. However, other morphological differences between the four species are also likely to be important in determining vibration transmission. Studies on other types of insect vibrations have shown that flexible plant stems attenuate vibrations more than stiff stems, as do thick leaves compared with thin leaves (Cocroft et al., 2006; Velilla et al., 2020). In buzz-pollinated flower, traits affecting vibration properties might include anther curvature (e.g. *S. houstonii*), stamen stiffness and length (Nunes et al., 2020). Similarly, the size of the anther locules and thickness of the anther walls may affect vibration transmission. Few studies have examined the effect of specific morphological traits on vibration transmission in buzz-pollinated flowers, but closely-related species of *Solanum* with similar morphologies differ in their vibration transmission properties (Arroyo-Correa et al., 2019). Moreover, the removal of stamen structures, such as the connective appendages in *Huberia bradeana* (Melastomataceae), can affect the relative amplitude of vibrations (Bochorny et al.).

Although our results only described the vibration properties of four species, the significant differences in the transmission of vibrations between taxa with different anther arrangements warrants further investigation. We hypothesise that the species with either “cone” or “loose” anther arrangements generally differ in their capacity to transmit vibrations, with implications for patterns of pollen release and interaction with buzz pollinators. If this is the case, bee pollinators may display different behavioural strategies to buzz these flowers and maximize pollen removal, for example, by changing the manipulation of anthers during visitation. For instance, we predict that bees on flowers with loose anther arrangements might learn to grab multiple anthers if this resulted in more efficient vibration transmission and thus a higher rate of pollen collection (Buchmann and Cane, 1989). Changes in floral handling might also influence the precision of pollen placement on bee bodies, and thus the effectiveness of “safe sites” (Koch et al., 2017; Vallejo-Marin et al., 2009). Further studies of how vibrations are applied to flowers with different androecium architectures and their effect on pollen release, including their placement on pollinators’ bodies, in both lab and field, will help ascertain the functional consequences of the enormous morphological diversity observed in buzz-pollinated flowers.

## Acknowledgements

We are grateful to Laurie Follweiler for greenhouse care. We acknowledge this work was performed on unceded traditional territory of the Osage.

## Funding

This work was supported by a NERC DTP-IAPETUS PhD studentship to LN, a Leverhulme Trust Research Grant to MVM (RPG-2018-235), and a Pittsburgh Ecology and Evolution Postdoctoral fellowship to A.L. Russell via the Dietrich School of Arts and Sciences.

## Data availability

Data will be deposited in the publicly available repository DataStorre.

---

**Supplementary Table 1.** Linear model for the x axis fitted with D_RMS_ as response, and forceps D_RMS_, anther type and species as fixed effects. \**P*-value of fixed effect in linear model. \*\**P*-value calculated using Type III sums of squares. Sample size is 150.

| X axis | Estimate | Std. error | P-value* | P-value** |
| --- | --- | --- | --- | --- |
| **Displacement amplitude D<sub>RMS</sub> (μm)** |  |  |  |  |
| Forceps D<sub>RMS</sub> | 0.946 | 0.056 | <0.001 | <0.001 |
| Anther (Non-focal) | -7.696 | 24.43 | 0.078 | 0.753 |
| Species |  |  |  | 0.846 |
| (*Exacum affine*) | 9.708 | 20.738 | 0.640 |  |
| (*Solanum dulcamara*) | 7.982 | 22.914 | 0.728 |  |
| (*Solanum houstonii*) | 18.722 | 21.106 | 0.887 |  |
| Forceps D<sub>RMS</sub>: Anther (non-focal) | -0.037 | 0.082 | 0.652 | 0.652 |
| Anther (non-focal): Species |  |  |  | <0.005 |
| Anther: (*Exacum affine*) | 29.011 | 29.623 | 0.329 |  |
| Anther: (*Solanum dulcamara*) | 14.929 | 32.161 | 0.643 |  |
| Anther: (*Solanum houstonii*) | 108.014 | 30.903 | <0.001 |  |

**Supplementary Table 2.**
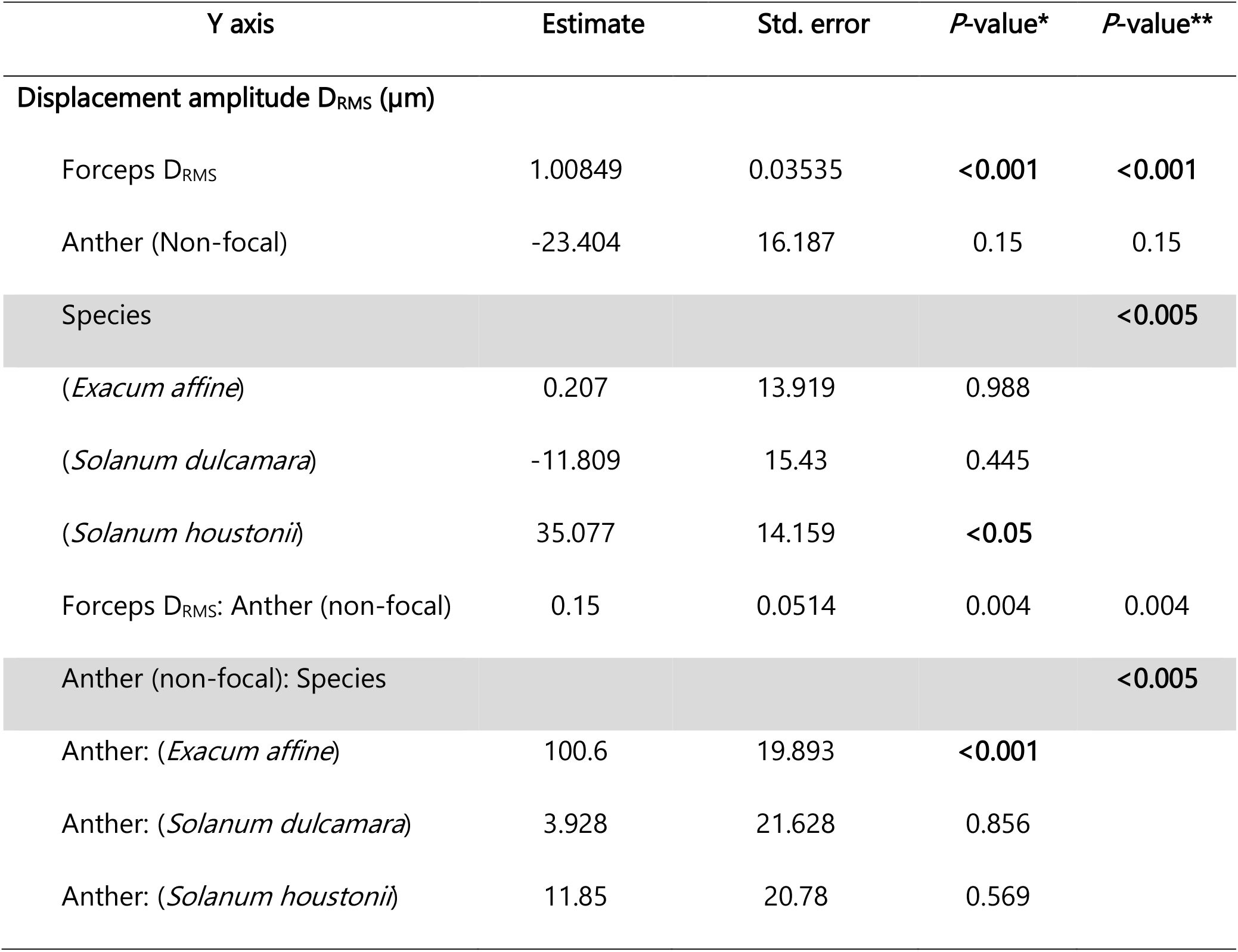
Linear model for the y axis fitted with D_RMS_ as response, and forceps D_RMS_, anther type and species as fixed effects. \**P*-value of fixed effect in linear model. \*\**P*-value calculated using Type III sums of squares. Sample size is 150.

| Y axis | Estimate | Std. error | P-value* | P-value** |
| --- | --- | --- | --- | --- |
| **Displacement amplitude D<sub>RMS</sub> (μm)** |  |  |  |  |
| Forceps D<sub>RMS</sub> | 1.00849 | 0.03535 | <0.001 | <0.001 |
| Anther (Non-focal) | -23.404 | 16.187 | 0.15 | 0.15 |
| Species |  |  |  | <0.005 |
| (*Exacum affine*) | 0.207 | 13.919 | 0.988 |  |
| (*Solanum dulcamara*) | -11.809 | 15.43 | 0.445 |  |
| (*Solanum houstonii*) | 35.077 | 14.159 | <0.05 |  |
| Forceps D<sub>RMS</sub>: Anther (non-focal) | 0.15 | 0.0514 | 0.004 | 0.004 |
| Anther (non-focal): Species |  |  |  | <0.005 |
| Anther: (*Exacum affine*) | 100.6 | 19.893 | <0.001 |  |
| Anther: (*Solanum dulcamara*) | 3.928 | 21.628 | 0.856 |  |
| Anther: (*Solanum houstonii*) | 11.85 | 20.78 | 0.569 |  |

**Supplementary figure 1.**
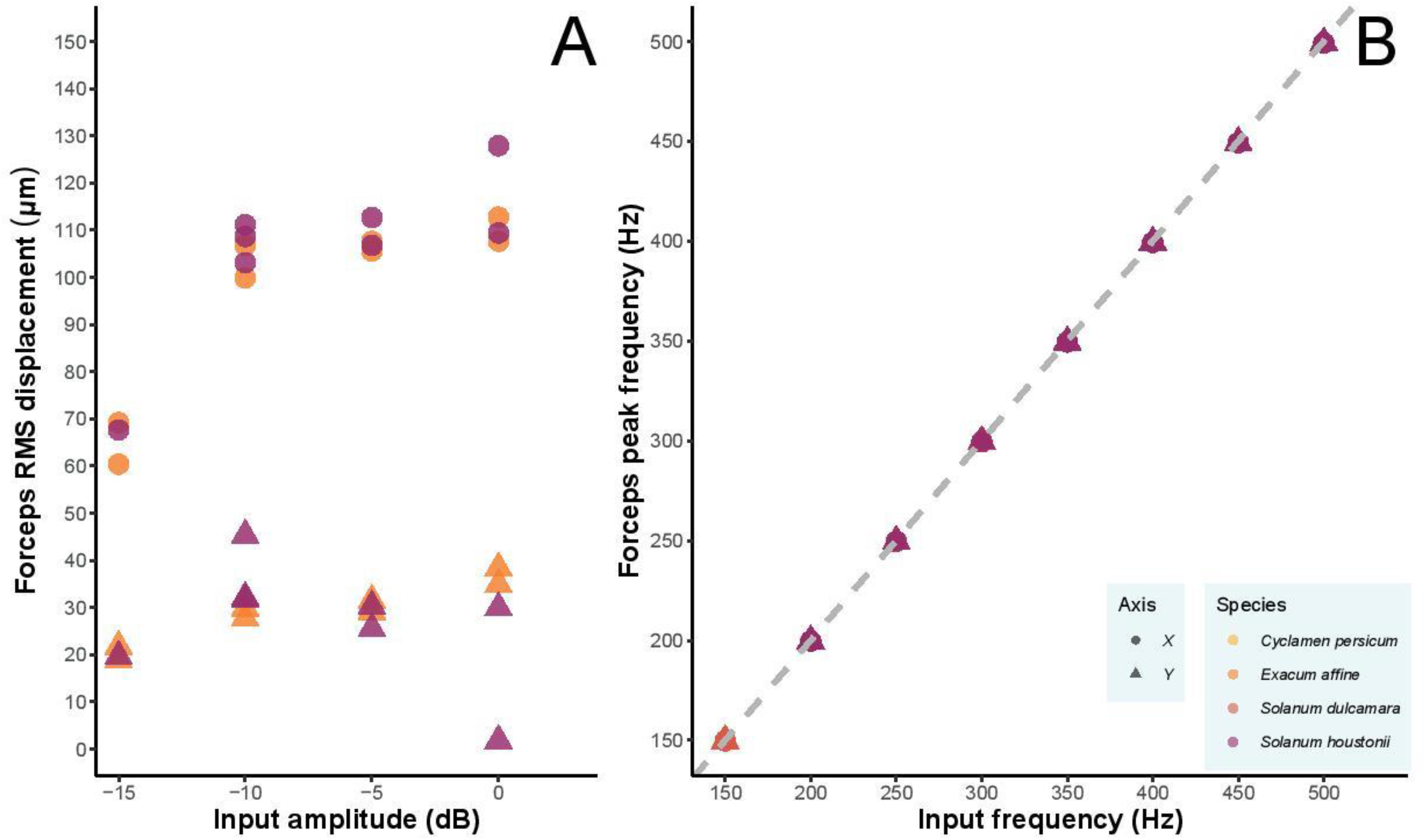
Measured RMS displacement (μm) of forceps against input amplitude (dB) (A) and measured peak frequency (Hz) of forceps against input frequency (Hz) (B). Grey dashed line indicates a linear relationship with slope=1.

**Supplementary figure 2.**
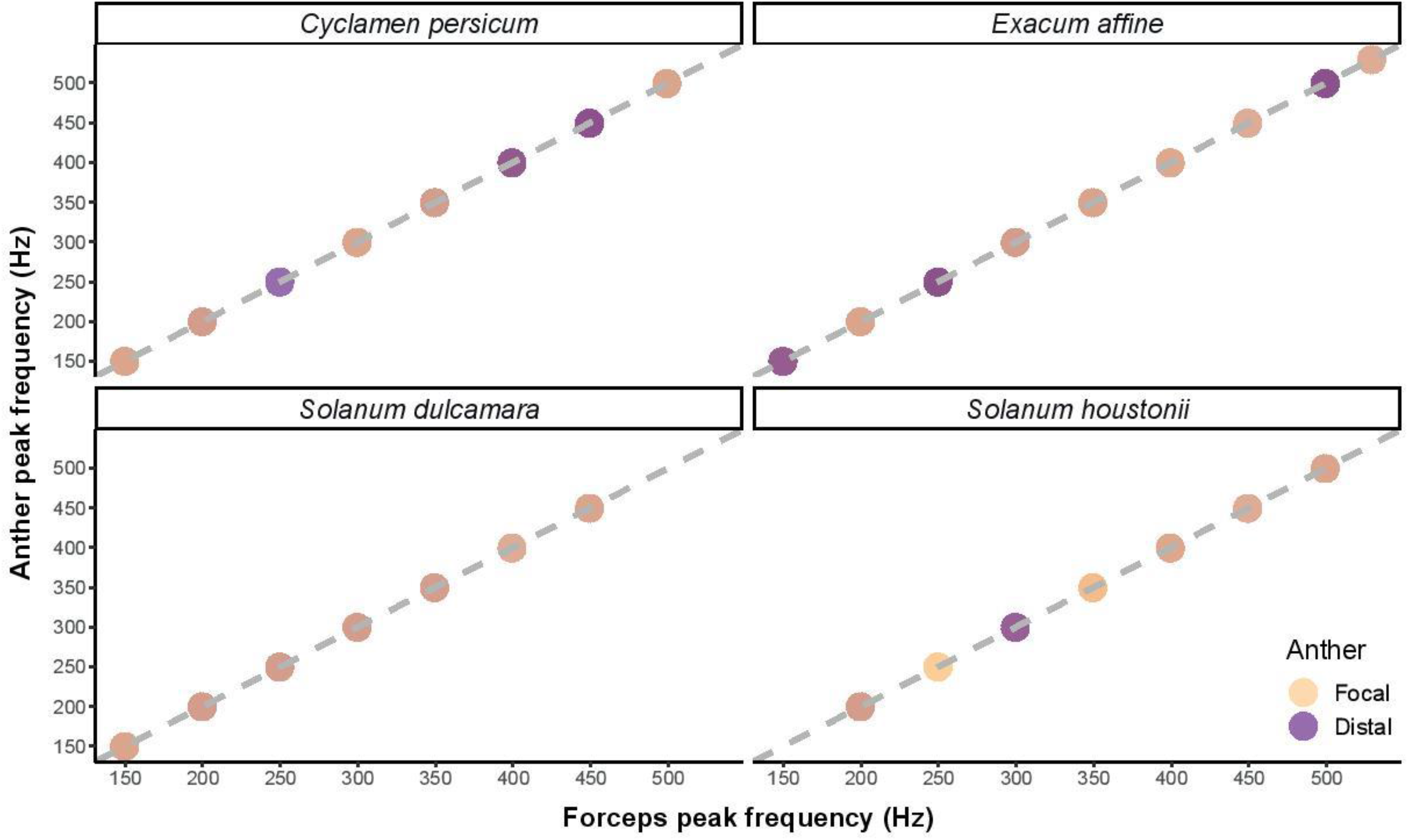
Measured peak frequency (Hz) against forceps frequency (Hz) for focal and distal anther of four plant species. Grey dashed line indicates a linear relationship with slope=1

**Supplementary Figure 3a.**
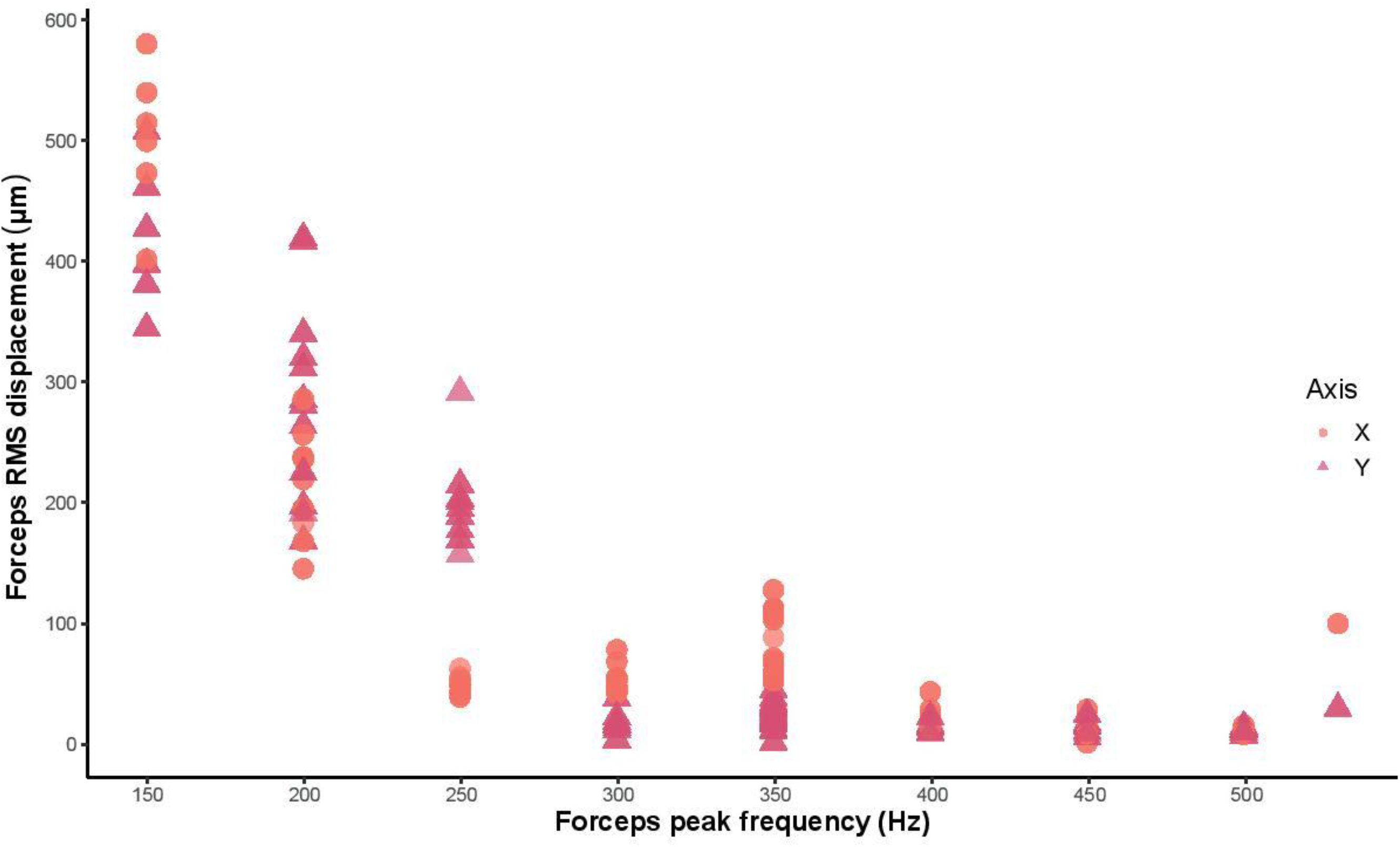
Forceps peak frequency (Hz) v forceps RMS displacement (μm) for both axes.

**Supplementary figure 3b.**
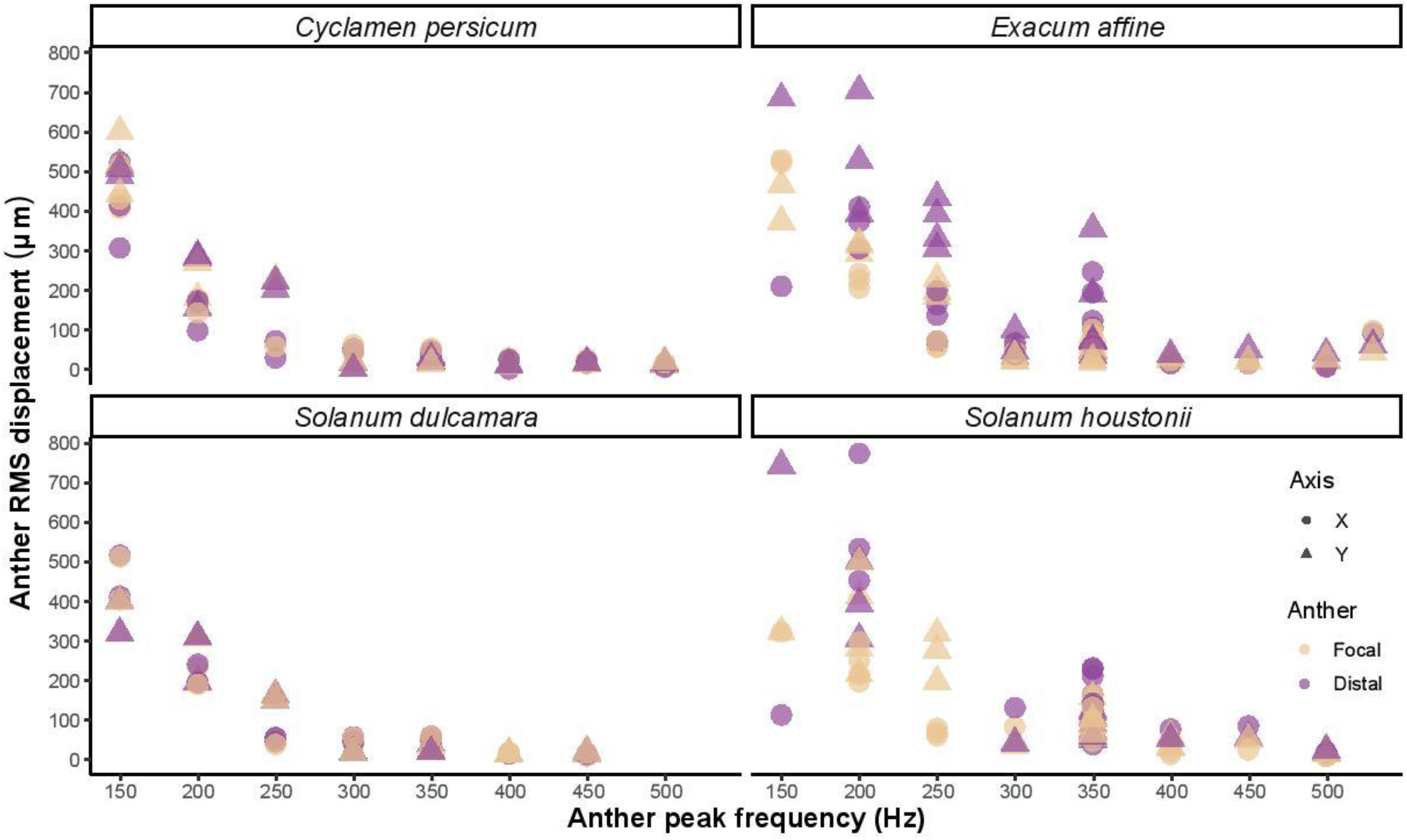
Anther peak frequency v anther RMS displacement (μm) for both axes.

**Supplementary Figure 4.**
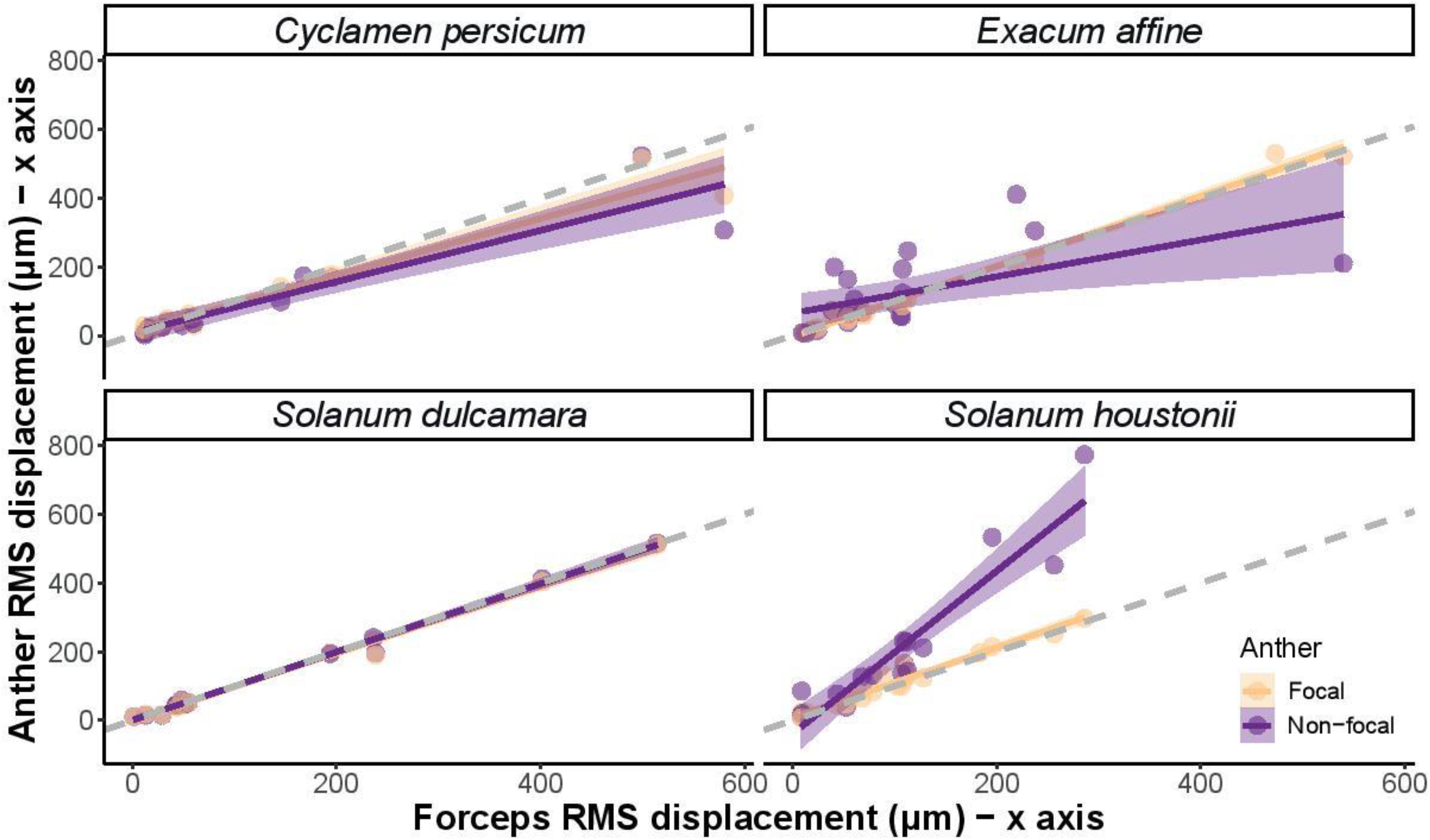
Linear model estimates and data points for measured x-axis RMS displacement (μm) of focal and distal anther against forceps RMS displacement (μm) in four plant species. Grey dashed line indicates a linear relationship with slope=1.

**Supplementary Figure 5.**
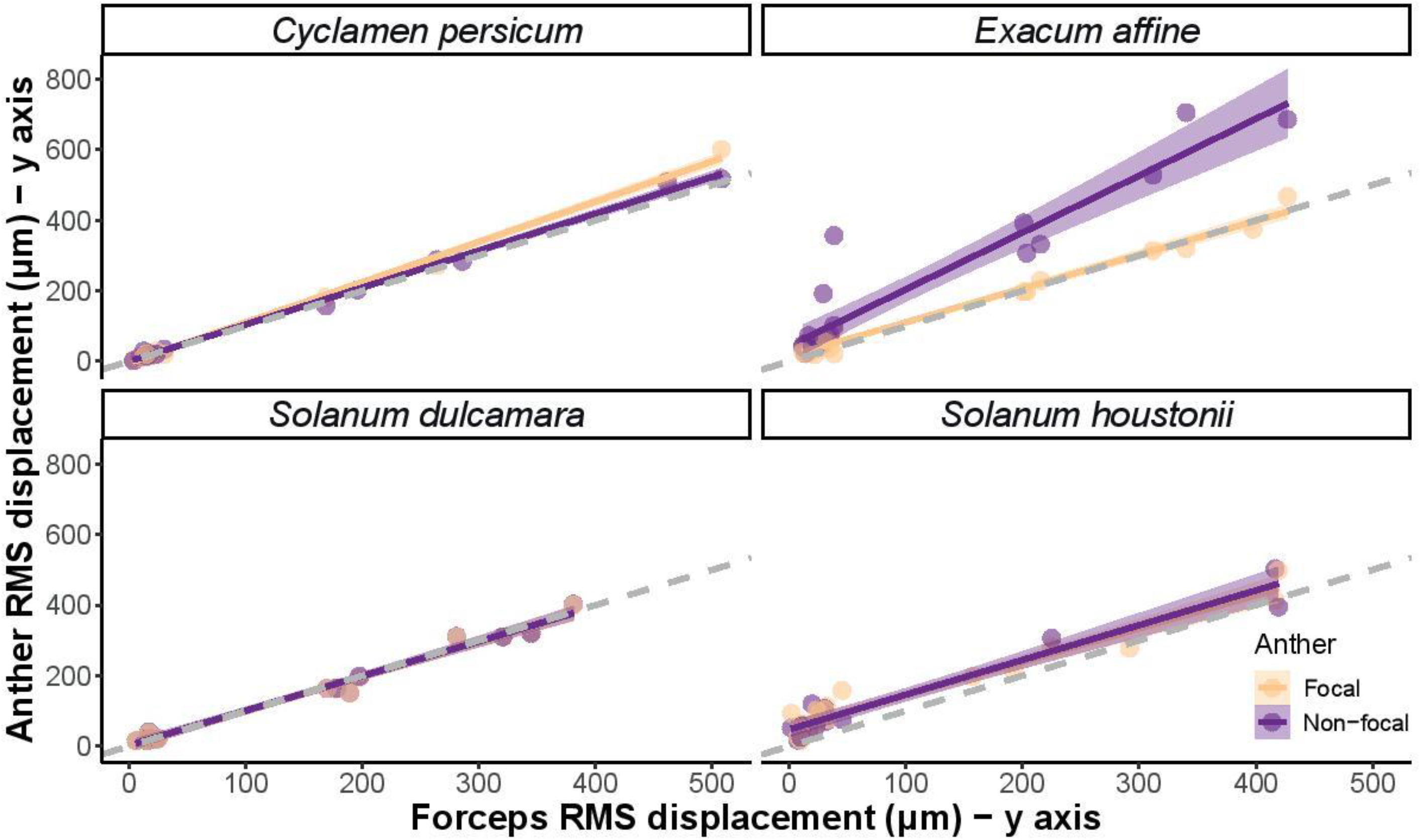
Linear model estimates and data points for measured y-axis RMS displacement (μm) of focal and distal anther against forceps RMS displacement (μm) in four plant species. Grey dashed line indicates a linear relationship with slope=1.

## Notes

### Competing Interest Statement

The authors have declared no competing interest.

### Summary of Updates

Text updated to refer to Figures S1 and S2.

## References

Arceo-Gómez, G., Martínez, M. L., Parra-Tabla, V. and García-Franco, J. G. (2011). Anther and stigma morphology in mirror-image flowers of *Chamaecrista chamaecristoides* (Fabaceae): implications for buzz pollination. Plant Biology 13 Suppl 1, 19–24.

Arroyo-Correa, B., Beattie, C. and Vallejo-Marín, M. (2019). Bee and floral traits affect the characteristics of the vibrations experienced by flowers during buzz pollination. Journal of Experimental Biology 222, jeb198176.

Bochorny, T., Bacci, L. F., Dellinger, A. S., Michelangeli, F. A., Goldenberg, R. and Brito, V. L. G. Connective appendages in *Huberia bradeana* (Melastomataceae) affect pollen release during buzz pollination. Plant Biology.

Brito, V. L. G., Nunes, C. E. P., Resende, C. R., Montealegre-Zapata, F. and Vallejo-Marín, M. (2020). Biomechanical properties of a buzz-pollinated flower. Royal Society Open Science 7, 201010.

Buchmann, S. L. (1983). Buzz pollination in angiosperms, pp. 73–113. New York, NY: Van Nostrand Reinhold Company.

Buchmann, S. L. (1985). Bees use vibration to aid pollen collection from non-poricidal flowers. Journal of the Kansas Entomological Society 58, 517–525.

Buchmann, S. L. and Cane, J. H. (1989). Bees assess pollen returns while sonicating *Solanum* flowers. Oecologia 81, 289–294.

Carbonell, A. K. Z. (2019). Expression and functional significance of andromonoecy in Solanum houstonii Martyn, vol. PhD thesis: University of Stirling, Stirling.

Cardinal, S., Buchmann, S. L. and Russell, A. L. (2018). The evolution of floral sonication, a pollen foraging behavior used by bees (Anthophila). Evolution 72, 590–600.

Cocroft, R. B. and Rodríguez, R. L. (2005). The behavioral ecology of insect vibrational communication. BioScience 55, 323–334.

Cocroft, R. B., Shugart, H. J., Konrad, K. T. and Tibbs, K. (2006). Variation in plant substrates and its consequences for insect vibrational communication. Ethology 112, 779–789.

Corbet, S. A. and Huang, S.-Q. (2014). Buzz pollination in eight bumblebee-pollinated *Pedicularis* species: does it involve vibration-induced triboelectric charging of pollen grains? Annals of Botany 114, 1665–1674.

De Luca, P. A., Buchmann, S., Galen, C., Mason, A. C. and Vallejo-Marín, M. (2019). Does body size predict the buzz-pollination frequencies used by bees? Ecology and Evolution 9, 4875–4887.

De Luca, P. A., Bussiere, L. F., Souto-Vilaros, D., Goulson, D., Mason, A. C. and Vallejo-Marín, M. (2013). Variability in bumblebee pollination buzzes affects the quantity of pollen released from flowers. Oecologia 172, 805–816.

De Luca, P. A., Giebink, N., Mason, A. C., Papaj, D. and Buchmann, S. L. (2020). How well do acoustic recordings characterize properties of bee (Anthophila) floral sonication vibrations? Bioacoustics 29, 1–14.

De Luca, P. A. and Vallejo-Marín, M. (2013). What’s the ‘buzz’ about? The ecology and evolutionary significance of buzz-pollination. Current Opinion in Plant Biology 16, 429–35.

Endress, P. (2012). The immense diversity of floral monosymmetry and asymmetry across angiosperms. The Botanical Review 78.

Faegri, K. (1986). The solanoid flower. Botanical Journal of Scotland 45, 51–59.

Free, J. B. (1970). The flower constancy of bumblebees. Journal of Animal Ecology 39, 395–402.

Glover, B. J., Bunnewell, S. and Martin, C. (2004). Convergent evolution within the genus *Solanum*: the specialised anther cone develops through alternative pathways. Gene 331, 1–7.

Harder, L. and Barclay, R. M. (1994). The functional significance of poricidal anthers and buzz pollination: controlled pollen removal from *Dodecatheon*. Functional Ecology 8, 509–517.

Hartig, F. (2019). DHARMa: residual diagnostics for hierarchical (multi-level/mixed) regression models. In R package version 0.2, vol. 4.

Hedrick, T. L. (2008). Software techniques for two- and three-dimensional kinematic measurements of biological and biomimetic systems. Bioinspir Biomim 3, 034001.

Kemp, J. E. and Vallejo-Marin, M. (2020). Pollen dispensing schedules in buzz-pollinated plants: Experimental comparison of species with contrasting floral morphologies. bioRxiv, 10.1101/2020.08.04.235739.

King, M. and Buchmann, S. (1996). Sonication dispensing of pollen from *Solanum laciniatum* flowers. Functional Ecology 10, 449–456.

King, M. and Buchmann, S. (2003). Floral sonication by bees: Mesosomal vibration by *Bombus* and *Xylocopa*, but not *Apis* (Hymenoptera: Apidae), ejects pollen from poricidal anthers. Journal of the Kansas Entomological Society 76, 295–305.

King, M. J. (1993). Buzz foraging mechanism of bumble bees. Journal of Apicultural Research 32, 41–49.

Koch, L., Lunau, K. and Wester, P. (2017). To be on the safe site – Ungroomed spots on the bee’s body and their importance for pollination. Plos One 12, e0182522.

Kollasch, A. M., Abdul-Kafi, A.-R., Body, M. J. A., Pinto, C. F., Appel, H. M. and Cocroft, R. B. (2020). Leaf vibrations produced by chewing provide a consistent acoustic target for plant recognition of herbivores. Oecologia 194, 1–13.

Ludecke, D. (2021). sjPlot - Data visualization for statistics in social science. In Zenodo.

Luo, Z., Zhang, D. and Renner, S. S. (2008). Why two kinds of stamens in buzz-pollinated flowers? Experimental support for Darwin’s division-of-labour hypothesis. Functional Ecology 22, 794–800.

Macior, L. W. (1964). An experimental study of the floral ecology of *Dodecatheon meadia*. American Journal of Botany 51, 96–108.

Mortimer, B. (2017). Biotremology: Do physical constraints limit the propagation of vibrational information? Animal Behaviour 130, 165–174.

Müller, H. (1881). Two kinds of stamens with different functions in the same flower. Nature 24, 307–308.

Nunes, C. E. P., Nevard, L., Montealegre-Zapata, F. and Vallejo-Marin, M. (2020). Are flowers tuned to buzzing pollinators? Variation in the natural frequency of stamens with different morphologies and its relationship to bee vibrations. bioRxiv, 2020.05.19.104422.

Oberst, S., Lai, J. C. S. and Evans, T. A. (2019). Physical basis of vibrational behaviour: channel properties, noise and excitation signal extraction. In Biotremology: Studying Vibrational Behavior, eds. P. S. M. Hill R. Lakes-Harlan V. Mazzoni P. M. Narins M. Virant-Doberlet and A. Wessel), pp. 53–78. Cham: Springer International Publishing.

Papaj, D. R., Buchmann, S. L. and Russell, A. L. (2017). Division of labor of anthers in heterantherous plants: flexibility of bee pollen collection behavior may serve to keep plants honest. Arthropod-Plant Interactions 11, 307–315.

Pritchard, D. J. and Vallejo-Marín, M. (2020). Floral vibrations by buzz-pollinating bees achieve higher frequency, velocity and acceleration than flight and defence vibrations. The Journal of Experimental Biology 223, jeb220541.

Puff, C., Igersheim, A., Buchner, R. and Rohrhofer, U. (1995). The united stamens of Rubiaceae: morphology, anatomy; their role in pollination ecology. Annals of the Missouri Botanical Garden 82, 357–382.

Rosi-Denadai, C. A., Araújo, P. C. S., Campos, L. A. d. O., Cosme Jr., L. and Guedes, R. N. C. (2020). Buzz-pollination in Neotropical bees: genus-dependent frequencies and lack of optimal frequency for pollen release. Insect Science 27, 133–142.

Russell, A. L., Golden, R. E., Leonard, A. S. and Papaj, D. R. (2015). Bees learn preferences for plant species that offer only pollen as a reward. Behavioral Ecology 27, 731–740.

Russell, A. L., Kikuchi, D. W., Giebink, N. W. and Papaj, D. R. (2020). Sensory bias and signal detection trade-offs maintain intersexual floral mimicry. Philos Trans R Soc Lond B Biol Sci 375, 20190469.

Schwartz-Tzachor, R., Dafni, A., Potts, S. and Eisikowitch, D. (2006). An ancient pollinator of a contemporary plant (*Cyclamen persicum*): When pollination syndromes break down. Flora 201, 370–373.

Sueur, J., Aubin, T. and Simonis, C. (2008). Seewave, a free modular tool for sound analysis and synthesis. Bioacoustics 18, 213–226.

Tong, Z.-Y., Wang, X.-P., Wu, L.-Y. and Huang, S.-Q. (2019). Nectar supplementation changes pollinator behaviour and pollination mode in *Pedicularis dichotoma*: implications for evolutionary transitions. Annals of Botany 123, 373–380.

Vallejo-Marín, M. (2019). Buzz pollination: studying bee vibrations on flowers. New Phytologist 224, 1068–1074.

Vallejo-Marín, M., Da Silva, E. M., Sargent, R. D. and Barrett, S. C. H. (2010). Trait correlates and functional significance of heteranthery in flowering plants. New Phytologist 188, 418–425.

Vallejo-Marin, M., Manson, J., Thomson, J. and Barrett, S. (2009). Division of labour within flowers: Heteranthery, a floral strategy to reconcile contrasting pollen fates. Journal of Evolutionary Biology 22, 828–39.

Vallejo-Marín, M. and Vallejo, G. C. (2021). Comparison of defence buzzes in hoverflies and buzz-pollinating bees. Journal of Zoology 3134, 237–249.

Varennes, L. P., Krapp, H. G. and Viollet, S. (2019). A novel setup for 3D chasing behavior analysis in free flying flies. J Neurosci Methods 321, 28–38.

Velilla, E., Polajnar, J., Virant-Doberlet, M., Commandeur, D., Simon, R., Cornelissen, J. H. C., Ellers, J. and Halfwerk, W. (2020). Variation in plant leaf traits affects transmission and detectability of herbivore vibrational cues. Ecology and Evolution 10, 12277–12289.

Vogel, S. (2013). Comparative Biomechanics: Princeton University Press.

Weisberg, S. and Fox, J. (2011). An R Companion to Applied Regression.

